# Combining behavior and mechanics approaches reveals the dynamics of animal impacts in mantis shrimp (Stomatopoda)

**DOI:** 10.1101/2023.11.20.567920

**Authors:** P.A. Green

## Abstract

Animals deliver and withstand physical impacts in diverse behavioral contexts, from competing rams clashing their antlers together to archerfish impacting prey with jets of water. Though the ability of animals to withstand impact has generally been studied by focusing on morphology, behaviors may also influence impact resistance. Mantis shrimp exchange high-force strikes on each other’s coiled, armored telsons (tailplates) during contests over territory. Prior work has shown that telson morphology has high impact resistance. I hypothesized that the behavior of coiling the telson also contributes to impact energy dissipation. By measuring impact dynamics from high-speed videos of strikes exchanged during contests between freely-moving animals, I found that over 20% more impact energy was dissipated as compared to a prior study that focused solely on morphology. This increase is likely due to behavior: because the telson is lifted off the substrate, the entire body flexes after contact, dissipating more energy than exoskeletal morphology does on its own. While variation in the degree of telson coil did not affect energy dissipation, higher velocity strikes resulted in greater energy dissipation, suggesting striking individuals may vary their behavior to affect impacts. Overall, these findings show that analysis of both behavior and morphology is crucial to understanding impact resistance, and suggest future research on the evolution of structure and function under the selective pressure of biological impacts.

**Summary statement:** Freely competing mantis shrimp dissipated over 90% of the energy of high-force strikes by raising their impact-resistant tailplates off the substrate; faster strikes led to greater energy dissipation.

## Introduction

Animals deliver and sustain physical impacts across contexts, from capturing prey (Vailati et al., 2012) to escaping predators (Patek et al., 2006). Impacts may be particularly prevalent in contests over limited and essential resources, where taxa as diverse as rams (Geist, 1966), parrotfish (Muñoz et al., 2012), and hermit crabs (Briffa and Elwood, 2001) collide body parts together in fights over resources. How impacts are received may determine injury costs and contest success. Through their importance to survival and reproduction, impacts may influence the evolution of animal morphology and behavior, selecting for materials, structures, and behaviors that help dissipate impact energy.

A significant body of literature has asked how materials and structures (hereafter combined as “morphology”, *sensu* Garland and Losos, 1994) withstand impacts. For example, techniques like finite element analysis (Drake et al., 2016; Johnson et al., 2021) and manipulative experiments (Kingston et al., 2022) have shown how bone and cuticle structures dissipate impact energy to protect animal brains. Other approaches have shown the importance of material properties to impact energy flow. For instance, trap-jaw ants close their jaws at ultrafast speeds, using these impacts for multiple purposes, from capturing soft-bodied prey to escaping predation by propelling themselves off the hard substrate (Spagna et al., 2009). By studying the kinematics of trap-jaw ant impacts, (Jorge et al., 2021) found that jaw closure transferred more energy to stiff targets than compliant ones, showing how the same impact, delivered to different materials, can achieve different biological tasks.

Impact energy dynamics may also be affected by behavior; that is, how animals use their morphology (*sensu* Garland and Losos, 1994). For example, hermit crabs compete over the shells they use to protect their bodies; during contests, one individual collides its shell with that of its opponent (Elwood and Briffa, 2001). These “attackers” vary in where they land impacts on the shell of a “defender” (Lane and Briffa, 2020) and the distance they displace their shell before starting their impact motion (Briffa and Fortescue, 2017). These behaviors could affect how impacts are felt by receivers. However, despite work in organismal biology that has emphasized the importance of both morphology and behavior to organismal performance (Garland and Losos, 1994; Green et al., 2021; Lauder, 1995), little research has taken this integrative approach to understand impact energy dissipation in animal systems.

Here, I use the coefficient of restitution (COR), a common engineering metric that quantifies the relative velocity of objects before and after their collision (Hibbeler, 1992), to understand the combined importance of behavior and morphology on impact energy dissipation. COR usually ranges from zero to one: a COR of zero represents a fully plastic impact, in which all impact energy is dissipated and the two colliding objects stick together, whereas a COR of one represents a fully elastic impact, such that no energy is lost and the objects bounce off in opposite directions with the same velocity as before their collision. Though COR is sometimes reported as a measure of a material (e.g., “the COR of a baseball bat is X”), it is more accurately a measure of a given impact (e.g., “the COR of a baseball bat hitting an 80mph curveball is X”), and factors such as the velocity and angle of impact can affect COR (Andersen et al., 1999; Chau et al., 2002; Haron and Ismail, 2012; Hibbeler, 1992).

I studied COR in mantis shrimp, an influential system in the study of biological impact (reviewed in deVries et al., 2021; Patek, 2019). During contests over burrow territories, mantis shrimp repeatedly deliver high-force impacts from raptorial appendages onto each other’s coiled telsons (tailplates) in a behavior termed “telson sparring” (Green and Patek, 2015). Both the appendage (Weaver et al., 2012) and telson (Yaraghi et al., 2019; Zhang et al., 2016) have morphological properties that help them withstand impacts, including increased mineralization at areas where contact occurs (Taylor and Patek, 2010).

Following methods common in the engineering literature, Taylor and colleagues (Taylor and Patek, 2010) measured the COR of the telson exoskeleton by dropping a steel ball onto a telson fixed to a lab bench and comparing the ball’s velocity post-impact to pre-impact. They found that the telson dissipated, on average, approximately 69% of the energy of an impacting strike (Taylor and Patek, 2010; Taylor et al., 2019). Further, COR varied with the body mass of the tested individual—the exoskeletons of larger shrimp dissipated more energy than those of smaller shrimp (Taylor and Patek, 2010).

In addition to the telson exoskeleton, how mantis shrimp use their telsons may affect impact energy dissipation during telson sparring. During natural interactions, instead of lying their telsons on the substrate, competitors raise their telsons in front of their bodies in a “telson coil” behavior (Fig. 1). The ability of the coiled telson to flex with the rest of the animal’s body after impact—much like a boxer moves with a punch he or she receives—might increase energy dissipation (see also Taylor and Patek, 2010). Furthermore, striking individuals can vary the angle and speed with which they strike in ways that may change energy dissipation. Green and colleagues (Green et al., 2019) found that mantis shrimp fighting against relatively smaller competitors delivered lower-velocity, lower-energy strikes than those fighting against relatively larger competitors. That is, competitors changed their striking behavior according to their opponent size. Variation in impact speed (Asteriou and Tsiambaos, 2018; Haron and Ismail, 2012) has been shown to lead to changes in energy dissipation in the engineering and sports science literature. Overall, in addition to morphological variation, behavioral variation might affect the COR of telson sparring impacts.

**Fig. 1.**
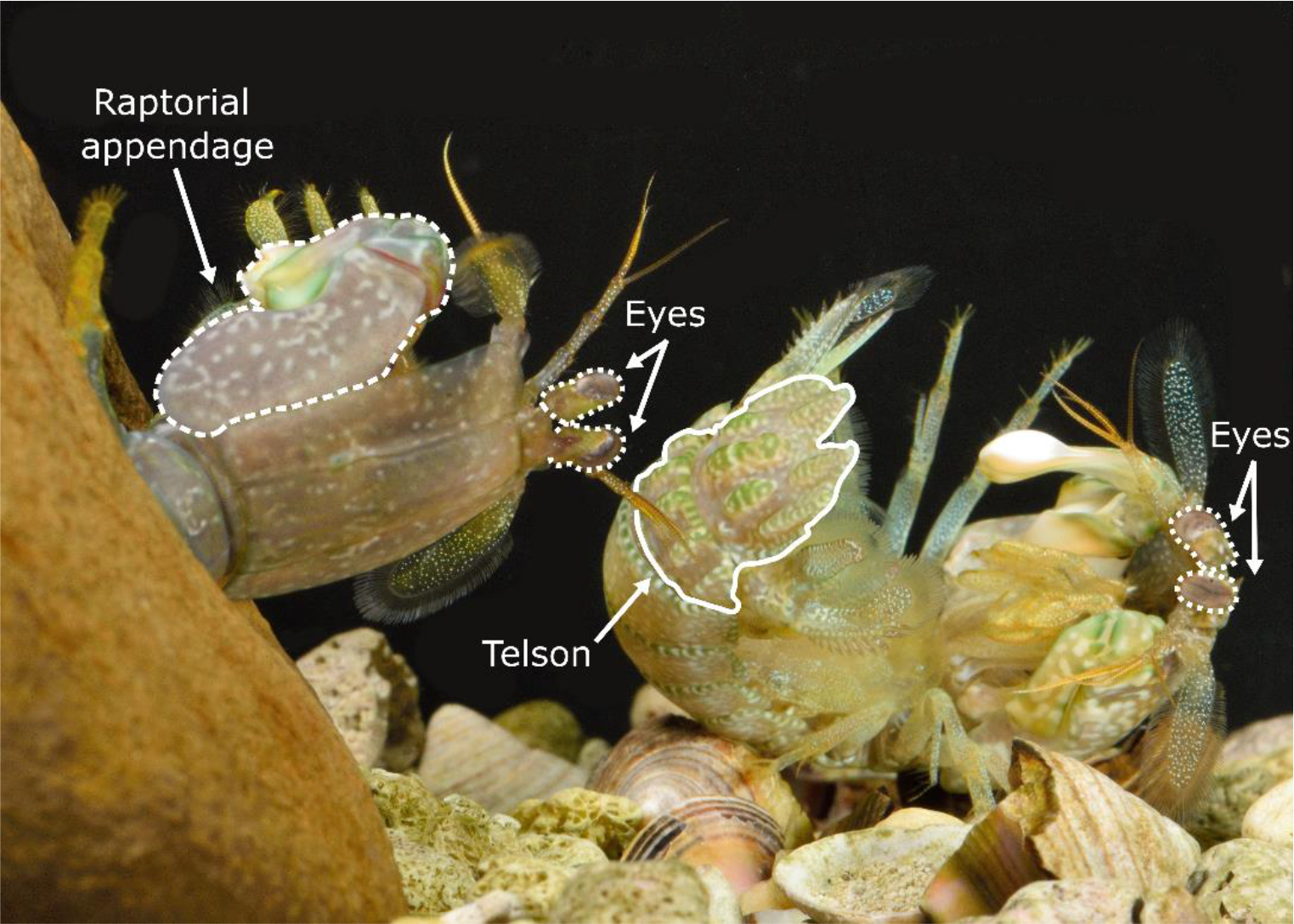
During contests, mantis shrimp lift their telsons off the substrate in a “telson coil” behavior. Here, the individual on the right is coiling its telson (solid white line) while lying on its left side (dorsal into page). The individual on the right is exiting a burrow (dorsal out of page), with its raptorial appendage outlined in dashed white. Both individual’s eyes are outlined in dotted white lines. Photo: Roy Caldwell.

I measured COR from live, sparring competitors. I first asked if and how my measures of COR that incorporated both morphology and behavior differed from those that focused only on morphology (Taylor and Patek, 2010; Taylor et al., 2019). I hypothesized that, because the telson and rest of the body can move freely *via* the telson coil behavior, the COR of impacts from live competitors would be lower (i.e., more energy dissipated) than that measured by (Taylor and Patek, 2010). I also asked whether variation in behavior or morphology that is both common to other studies of COR (impact velocity, object mass; e.g., Chau et al., 2002; Cross, 2002; Haron and Ismail, 2012) and unique to the mantis shrimp system (angle of the telson coil, angle of the appendage at contact) affected variation in COR. Following the engineering literature, I hypothesized that greater appendage mass of striking individuals and greater impact velocity would result in decreased COR (Asteriou and Tsiambaos, 2018; Chau et al., 2002; Haron and Ismail, 2012). I also hypothesized that, if the telson coil behavior has evolved (at least in part) to facilitate impact energy dissipation, increases in the degree to which the telson was coiled would result in lower COR. Finally, I tested the effect of the angle of the appendage at contact (compared to before the strike motion) on COR; I did not have a directional hypothesis as to how appendage angle at contact would affect COR.

## Methods

The contests studied here were originally studied in (Green et al., 2019).

### Animal collection and care, and contest staging

I collected *N. bredini* from burrows in coral rubble in Panama (Autoridad Nacional del Ambiente collection permits SE/A-115-13; SE/A-92-15; SE/A-52-17), transported them to Duke University, and housed them individually in 10cm^3^ plastic cubes in an aquarium with circulating artificial sea water (27° C, 12h:12h light:dark cycle). Each individual was provided a burrow refuge made of PVC tubing cut longitudinally and secured to the side of the cube, and was fed twice weekly with frozen krill and brine shrimp, or fresh snails.

I staged contests between competitors that were matched for body size within the range of prior studies (Green and Patek, 2015; Green and Patek, 2018). Measurements of body mass and body length were taken after contests using a digital scale (Denver Instruments APX-3202 balance, Sartorius AG, Goettingen, Germany) and digital calipers (Mitutoyo Digimatic Caliper, Mitutoyo Corp., Kawasaki, Japan).

Contests were staged by introducing a competitor in a PVC burrow in front of the burrow of a second individual. After initially separating competitors with an opaque barrier, I removed the barrier and filmed sparring strikes with 30,000-40,000 frames/second high-speed video. Competitors readily sparred after the barrier was removed; fighting behaviors were similar to those observed in previously-studied interactions (Green and Patek, 2018).

### Quantifying appendage and telson motion, and COR

From high-speed videos, I digitized the movement of the appendage and the telson using landmarks or natural brightness patterns, following methods in (Kagaya and Patek, 2016) and using the MtrackJ plugin in Fiji v.1.53 (Meijering et al., 2012; Schindelin et al., 2012). I digitized appendage movement from the beginning of the strike motion until 10 frames after it made contact with the telson, and telson movement from 10 frames before the appendage made contact until 10 frames after contact (Fig. 2). Note that, in some cases, contact occurred between two frames; in these cases, I chose the latter frame as the frame of contact, to ensure I was identifying when contact occurred and not before.

**Fig. 2.**
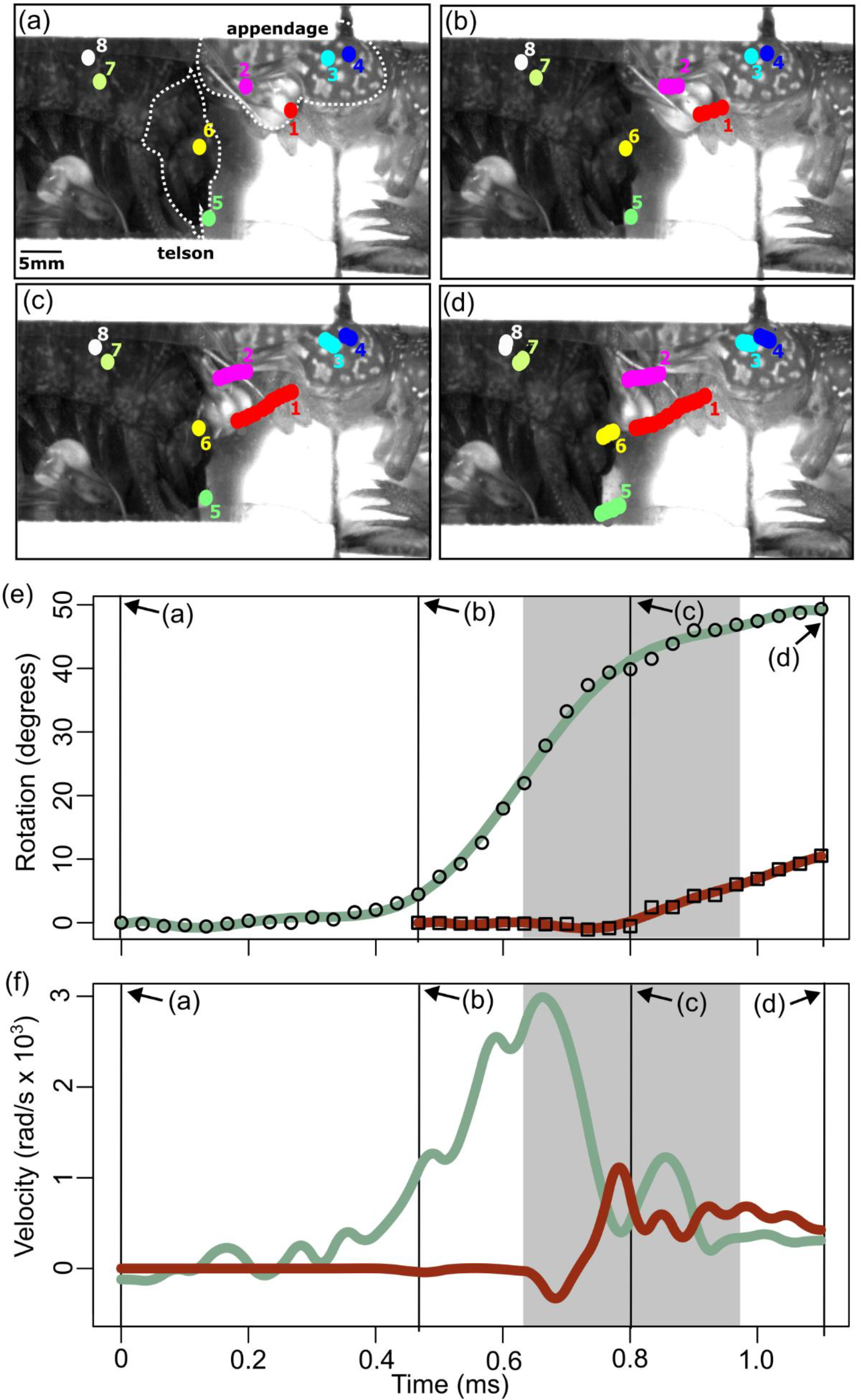
High-speed videos of strikes were used to quantify appendage and telson displacement and velocity, and to calculate COR. (a-d) Still images from high-speed video showing points digitized on striking body (1, 2) and merus (3, 4) of striking individual, and on telson (5, 6) and abdomen (7, 8) of individual receiving strike. Appendage and telson are outlined with dotted white lines and indicated in (a); individual receiving strike has appendage at bottom. (e) Plot of rotational displacement (y-axis) over time (x-axis) of raw data (points) and polynomial fit (lines) of appendage (open circles, green line) and telson (open squares, red line). (f) Plot of velocity (y-axis) over time (x-axis) of appendage (green line) and telson (red line). In both (e) and (f), solid vertical lines correspond to images in (a-d); line at 0.8ms is frame of contact. Shaded gray area indicates five frames before and after frame of contact, the range over which velocity (slope of displacement data) before and after contact was averaged.

From the digitized points, I generated rotational displacement data (Fig. 2e) using R (v. 4.2.1, Core Team, 2020) code originally published by Kagaya and Patek (2016) (Fig. 2e). I then used MatLab (R2023a) code developed in prior work (Green et al., 2019; McHenry et al., 2012; McHenry et al., 2016) to calculate appendage and telson velocity (Fig. 2f). This code fits a 7^th^-9^th^ order polynomial spline (spline order evaluated visually for each strike) to the raw displacement data, interpolates 5000 points along the spline, and takes the derivative of displacement with respect to time to calculate velocity.

From velocity data, I calculated COR using two methods. The first method replicates the ball-drop test used by Taylor and Patek (2010) and Taylor et al., (2019) by assuming the telson acts as a surface. This method calculates COR as:

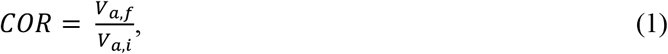

where *V*_*a,f*_ is the mean velocity of the appendage over five high-speed video frames after contact and *V*_*a,i*_ is the mean velocity of the appendage over five frames before contact.

The second method treats the telson as a second particle, similar to two billiard balls colliding, and accounting for the telson’s movement before and after contact (Hibbeler, 1992):

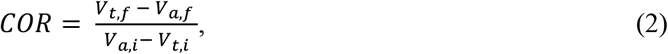

where *V*_*t,f*_ and *V*_*a,f*_ represent telson and appendage velocity after contact, respectively, and *V*_*t,i*_ and *V*_*a,i*_ represent telson and appendage velocity before contact, respectively. As in equation (1), velocity was the mean velocity across five frames before or after contact.

### Morphological and behavioral correlates of COR

I quantified hypothesized morphological and behavioral correlates of COR (see Introduction). These included (1) the angle of appendage displacement at the time of contact; (2) the velocity of the strike at the frame at which contact occurred, (3) the angle of the telson coil adopted by the individual receiving the strike; (4) the mass of the moving segments (dactyl and propodus) of the striking individual’s appendage, termed “striking body mass” (see also Green et al., 2019); and (5) whether or not the strike was a glancing blow. Metrics 1 and 2, the angle and velocity of the appendage at contact, were quantified through the MatLab code described above. Metric 3, the angle of the telson coil, was quantified as the interior angle of an open triangle drawn (in Fiji v 1.53) from the tip of the telson, to the distal point of the 5^th^ abdominal tergite (approximately halfway between the telson and carapace), to the distal edge of the carapace; lower values represented an individual that adopted a more coiled posture (Fig. 3 and Supplementary Video). Metric 4, the mass of the striking body, was quantified using a scaling relationship between striking body length and mass developed by Green et al. (2019). Metric 5 was a binary (yes/no) variable that was quantified by observing the high-speed videos (see examples in Supplementary Video).

**Fig. 3.**
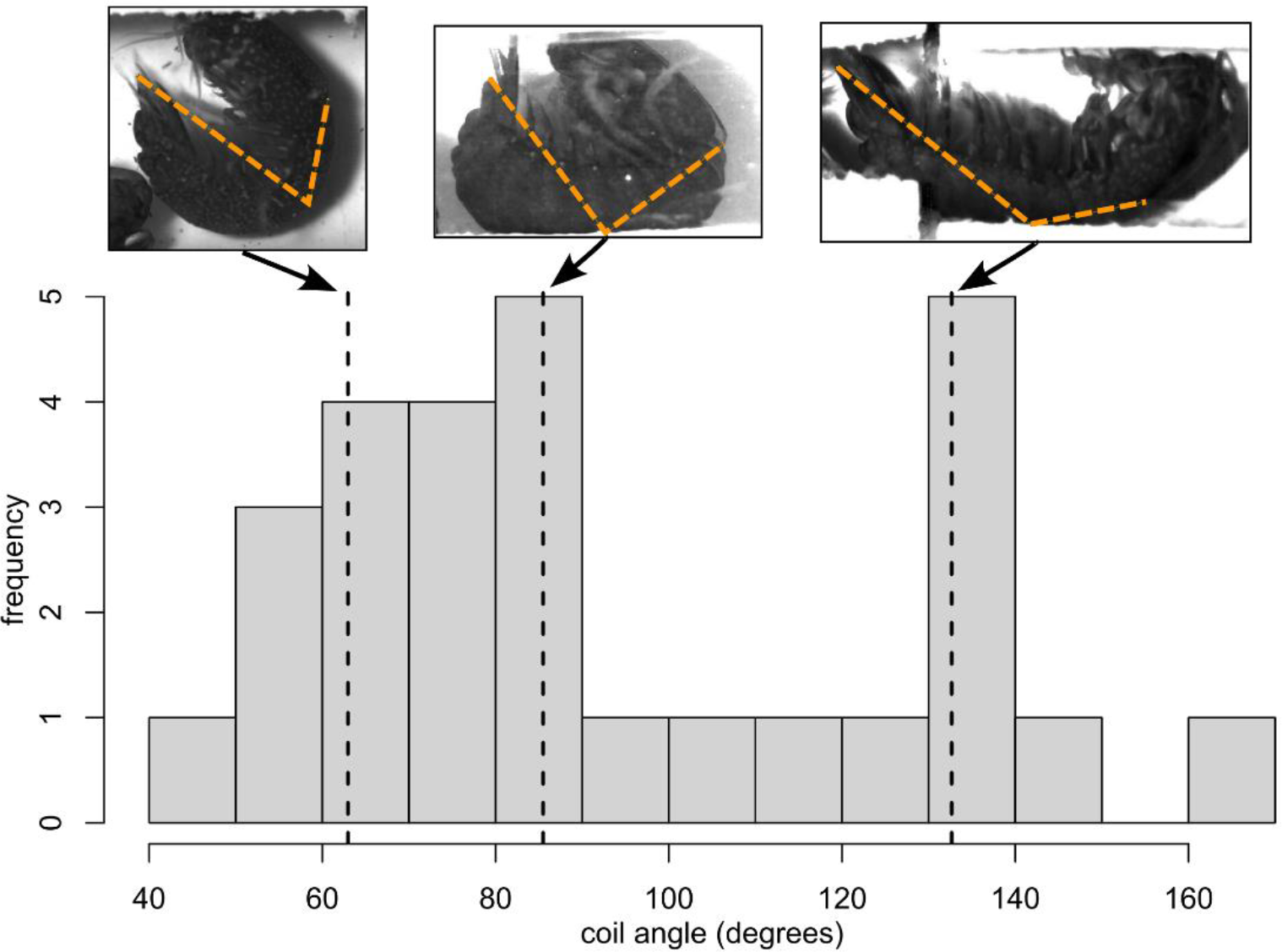
Histogram of variation in contact angle. Inset images above histogram correspond to dashed vertical lines, showing examples of contact angle variation. Dashed, orange lines on inset images show the lines used to calculate coil angle. In all example images, the dorsal side of the shrimp faces down.

### Statistical analyses

All statistical analyses were conducted in R v. 4.2.1 (Core Team, 2020).

I first compared measurements of COR to the raw data of Taylor and Patek (2010), found in the supporting data of Taylor et al. (2019). While these data were collected from *N. wennerae*, I filmed interactions between *N. bredini*. However, these species are closely related (Taylor et al., 2019); both Caribbean; and visually indistinguishable, including in telson morphology (*personal observation*). Further, measures of COR from a limited sample (N = 5 individuals) of *N. bredini* are within the range of the larger sample size (N = 16) of *N. wennerae* used here (Taylor et al., 2019).

I built a linear model predicting COR from data type: data from Taylor and Patek (2010), data from the present study using eqn. (1), and data from the present study using eqn. (2). After checking a histogram of model residuals to ensure normality, I tested for the significance of data type using the Anova function in car package (Fox and Weisberg, 2019), and conduced Tukey post-hoc tests of two-way comparisons using the emmeans function in emmeans package (Lenth, 2022).

To test what behavioral and morphological variables predicted variation in COR, I built two linear mixed models (lme4 package Bates et al., 2015) predicting COR as measured by eqn. (1) and, separately, eqn. (2). Predictor variables were those described above. I also included an interaction between strike velocity at contact and striking body mass. I initially included an interaction between appendage angle at contact and coil angle, but removed this due to high variance inflation factors (Zuur and Ieno, 2016; Zuur et al., 2010). Finally, I included random effects of the identity of both competitors to account for multiple strikes from some individuals. I checked histograms of model residuals to ensure good model fit and tested for significance on the full model using the Anova function in the car package.

## Results

I filmed 29 strikes from 10 unique striking individuals and 12 unique individuals receiving strikes. I gathered data on COR from appendage movement only (eqn. (1)) on all 29 strikes, and from appendage and telson movement (eqn. (2)) on 20 strikes. In nine cases, I could not digitize telson movement due to blocking of landmarks by cavitation bubbles or other obstructions.

Measures of COR from live sparring interactions were significantly lower, indicating more energy was dissipated, than those that measured morphology only (Fig. 4A; Table S1). Incorporating both appendage and telson motion (eqn. (2)) resulted in lower COR than measures of just appendage motion (eqn. (1); Fig 4A; Table S1). Notably, the mean COR when measured from both appendage and telson motion was negative. Negative COR values occurred when the appendage moved faster than the telson after contact (see, e.g., Fig. 2 Fig. S1). In these cases, the numerator of eqn. (2) (post-contact motion) was negative, whereas the denominator was always positive, as the appendage always moved faster than the telson pre-contact.

**Fig. 4.**
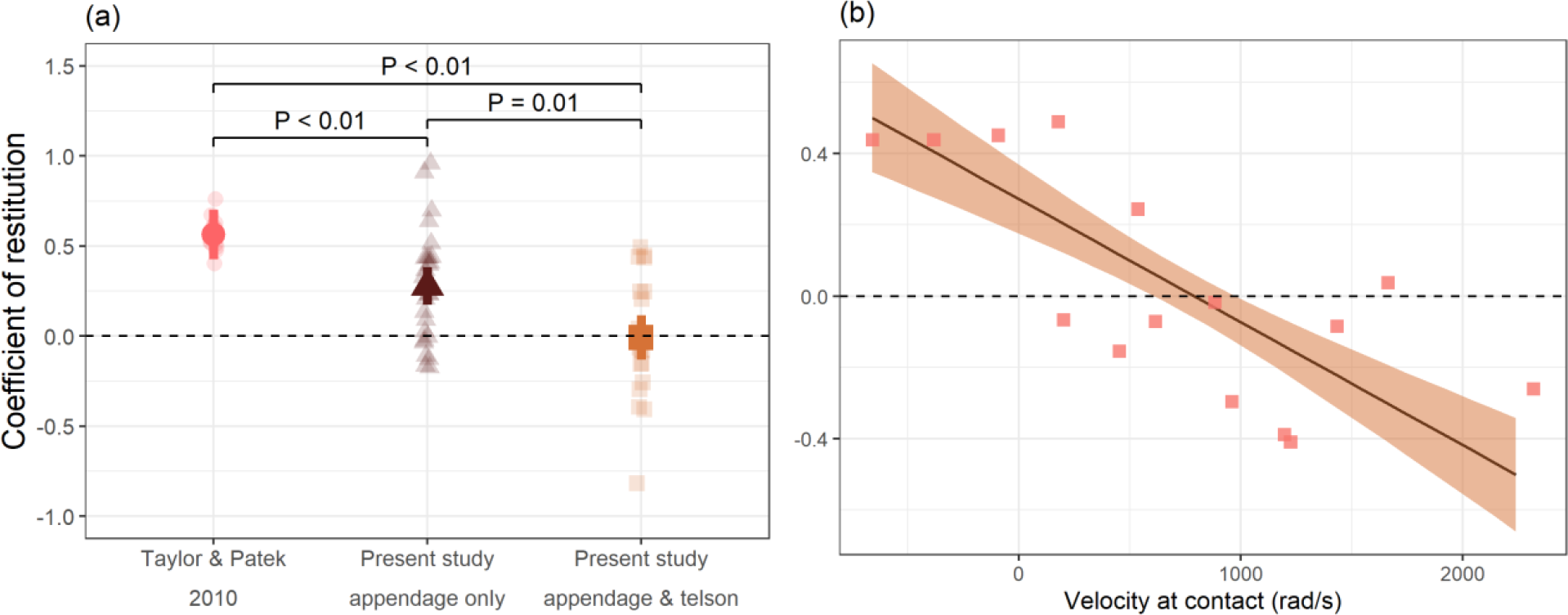
(a) COR was lower when incorporating both behavior and morphology. Pink circles represent data from Taylor and Patek (2010) (morphology only), brown triangles represent data from present study measuring appendage movement only, and orange squares represent data from present study measuring both appendage and telson movement. Semi-transparent points show raw data; solid points, error bars, brackets and text show estimates, 95% CIs, and P-values, respectively, from model described in main text. Sample sizes in (a) were: N = 16 strikes (Taylor and Patek, 2010); N = 29 strikes, 10 striking individuals, 12 struck individuals (present study, appendage movement only); N = 20 strikes, 9 striking individuals, 11 struck individuals (present study, appendage and telson movement). (b) COR for appendage and telson data (y-axis) decreased with increasing velocity at contact (x-axis; radians/second). Points represent raw data; solid line and ribbon represent estimate and standard error, respectively, from LMM described in text. Sample size in (b) was N = 15 strikes, 7 striking individuals, 7 struck individuals. In both (a) and (b), horizontal dashed line shows COR of zero.

Strikes with higher contact velocities resulted in lower COR (Fig. 4B); no other variables significantly correlated with COR (Table S2). This result held only when measuring appendage and telson motion (eqn. (2)); no measured variables significantly predicted energy dissipation when accounting for only appendage motion (eqn. (1); Table S3).

## Discussion

Measurements from freely interacting competitors reveal the contributions of both morphology and behavior to energy dissipation in animal impacts. Previous work using a ball-drop technique that focused only on the morphology of the telson exoskeleton found it dissipates approximately 69% of impact energy (energy dissipation calculated as 1 – COR^2^; Taylor and Patek, 2010). When comparing my data using the same equation as that used in a ball-drop test (eqn. (1)), I found mean energy dissipation of approximately 92%. This increase is most likely due to the telson coil; that is, the behavior with which competitors use their telson. By lifting the telson off the substrate, the whole body of the animal flexes upon receiving a strike (Supplementary Video), much like a boxer moving with a punch he or she receives. As a result, energy is dissipated throughout the body and not just in the telson exoskeleton, an effect that Taylor and Patek (2010) suggested in their prior study. Importantly, my results do not suggest those of Taylor and Patek (2010) are inaccurate. Rather, they add information to this prior study of morphology by showing how the behavior of the telson coil can affect energy dissipation. Other factors may also increase energy dissipation, including loss of energy to cavitation (Patek and Caldwell, 2005). I observed cavitation upon contact in every strike I recorded, and at times cavitation bubbles developed far beyond the location of the strike (e.g., on the ventral side of the telson or proximally along the abdomen).

Although COR is usually between 0 and 1 (Hibbeler, 1992), the mean COR when incorporating both appendage and telson motion (eqn. (2)) was negative (Fig. 4a). Negative COR values have also been described in engineering (Müller et al., 2012) and sports science (Cross, 2002), especially for oblique impacts like those studied here. More relevant to the biology of the mantis shrimp system, negative values resulting from high post-contact appendage velocity suggest the appendage may still be powered after the strike. This power could come from continued uncoiling of the appendage exoskeletal spring and/or from continued muscle contraction after contact. While mantis shrimp have been shown to vary strike kinematics according to behavioral context (fighting versus feeding, Green et al., 2019) and the medium in which they strike (water versus air, Feller et al., 2020), prior work in muscle physiology has suggested that kinematic variation is driven solely by variable muscle activation *before* the strike is released (e.g., Kagaya and Patek, 2016). Future work might look for muscle activity later in the strike motion, testing whether individuals “punch through” a target.

COR decreased with increasing appendage velocity at contact (Fig. 4b), a relationship similar to that found in engineering (Asteriou and Tsiambaos, 2018; Chau et al., 2002) and sports science (Haron and Ismail, 2012). This relationship only occurred when considering both telson and appendage movement, not appendage movement on its own, reflecting the importance of measurement technique to COR. When combining telson and appendage motion, greater appendage velocities after contact resulted in negative COR (see Results and eqn. (2)). By contrast, when considering appendage movement on its own, greater appendage velocities post-contact resulted in a more positive numerator (*V*_*f*_) of eqn. (1), leading to positive values of COR. Given that live animals can move freely, it seems more accurate to consider the telson as a second particle and not a stable surface. However, some aspects important to treating the telson as a second particle were beyond the scope of the present study. For example, analyses of two-particle impacts are improved by incorporating the angle of their collision (Hibbeler, 1992). While I measured the angle of appendage displacement at contact, I could not accurately measure the angle of contact between the appendage against the telson. Future work, for example using 3D videography approaches or mounting dead animals in coiled postures and conducting ball drop tests from multiple angles (e.g., Chau et al., 2002; Cross, 2002), might better model telson-appendage impacts.

One interpretation of the relationship between appendage velocity at contact and COR (Fig. 4b) is that striking individuals increase strike velocity to increase energy dissipation by their opponents that receive strikes. If strike energy dissipation is an active process by strike receivers—for example, if energy dissipation required muscle activation—then dissipating more energy from higher-velocity strikes might lead to more rapid exhaustion by strike receivers, leading to a more rapid retreat. This hypothesis could be tested by combining high-speed video approaches like those used here with measurements of muscle physiology or metabolism after contests. In hermit crabs, competitors that received more powerful impacts from their opponent’s shell (as measured by sound intensity) had decreased hemolymph glucose concentrations and were more likely to lose the contest (Briffa and Elwood, 2002); however, these physiological metrics were not linked to impact kinematics *per se*.

Outside of velocity at contact, no other factors I analyzed affected COR. For example, while it seems intuitive that strike receivers that coil their telson more tightly might dissipate more strike energy, the degree to which the telson was coiled did not predict COR. It could be that, once the telson is lifted off the substrate and the body can flex upon receiving a strike, changing the degree of telson coil has minimal importance. Indeed, although mantis shrimp dissipated more energy from higher-velocity strikes (Fig. 4b), at minimum they dissipated 76% of strike energy. Variation greater than this already-high value may have little importance, leading to relaxed selection on variation in the telson coil behavior. An alternative hypothesis for why individuals vary in the degree of telson coil relates to contest behaviors.

Individuals outside of burrows frequently coil their telsons tightly in front of their bodies (Fig. 1); this allows them to both receive strikes safely and rapidly uncoil to deliver strikes in return (Green and Patek, 2018). By contrast, individuals already in burrows often simply hold their telsons outside the burrow to receive strikes without tightly coiling them, receiving multiple strikes without delivering strikes themselves (personal observation). Coiling may therefore be more of a behavioral strategy for moving quickly from defense to attack, as opposed to a mechanism of increasing energy dissipation.

Finally, Taylor and Patek (2010) found that larger mantis shrimp (measured by total body mass) had telsons that dissipated more strike energy (i.e., lower COR values). I was unable to include the body mass of individuals receiving strikes in my models predicting COR because this variable was highly collinear with the mass of the striking individual’s striking body (Pearson correlation = 0.91). However, in follow-up models that replaced appendage striking body mass with the body size (total body mass) of the struck individual, I found no effect of size on COR (Supplemental Tables S4, S5). The key difference between my results and those of Taylor and Patek (2010) is that I matched opponents for body size. This was done to encourage individuals to strike, as size-matched opponents exchange more strikes (Green and Patek, 2018). As a result of this size matching, the relative mass of the striking object (the appendage striking body) and the target (strike receiver body) and were somewhat similar across strikes and individuals. In comparison, Taylor and Patek (2010) kept the mass of the striking object—a steel ball— consistent, while varying the mass of the target. Therefore, Taylor and Patek (2010) achieved greater variation in the relative object mass to target mass (coefficient of variation of Taylor and Patek (2010) object to target mass = 72.1; CV of present study object to target mass = 15.3). To better test if COR scales with body size in natural contests, future work could less closely match the size of sparring competitors.

Both classic (Garland and Losos, 1994; Lauder, 1995) and recent studies (Bauer et al., 2020; Green et al., 2021) in integrative organismal biology have discussed the importance of couching biomechanics in greater ecological or behavioral realism. Using engineering-based techniques to analyze impacts from freely-behaving competitors, my data suggest that both morphology and behavior—how morphology is used (*sensu* Garland and Losos, 1994)—contribute to impact resistance in mantis shrimp. Given the widespread nature of impacts in contexts from foraging (Spagna et al., 2009; Vailati et al., 2012) to competition (Elwood and Briffa, 2001; Muñoz et al., 2012), and the importance of impact resistance to survival and reproduction, incorporating analyses of morphology and behavior to understand impact energy dissipation could help us understand how impacts act as a selective force on organismal structure and function.

## Acknowledgments

Arjun Bhatt and Brooke Sauer helped with filming and digitizing strike movement. Kacey Rinehart helped with digitizing telson movement. Three anonymous reviewers and Drs. Philip Anderson, Eleanor Caves, Sönke Johnsen, Matt McHenry, Sheila Patek, and Jennifer Taylor provided valuable feedback.

## Competing Interests Statement

I have no competing interests.

## Funding

I was funded by Human Frontier Science Program Fellowship LT000460/2019-L; funding from UC Santa Barbara; and a Duke University Biology Fellowship, Summer Research Fellowship, and Grants in Aid.

Other funding included National Science Foundation grants to Sheila Patek (IOS 1439850) Matt McHenry (IOS 1354842).

## Data Availability

Datasets and relevant code are available on FigShare: https://figshare.com/s/18e9c700a87e696a927f. A DOI will be generated upon publication.

## Notes

### Competing Interest Statement

The authors have declared no competing interest.

